# Early evolution of the prokaryotic transcription factor repertoire

**DOI:** 10.64898/2026.04.08.717362

**Authors:** Inder Raj Singh, Akshara Dubey, Aswin Sai Narain Seshasayee

**Affiliations:** National Centre for Biological Sciences, Tata Institute of Fundamental Research, Bellary Road, Bangalore 560065, India; Manipal Academy of Higher Education, Manipal, India

## Abstract

Transcription initiation is regulated by proteins called transcription factors (TFs). Though TFs help determine phenotype across the tree of life, they are nonessential for minimal cellular life and are often absent in endosymbiotic and parasitic organisms. Given this and the idea that it is a certain level of organism ‘*complexity*’ that calls for specific transcription regulation, we traced the evolutionary history of TF repertoire on a bacterio-archaeal tree of life using a dataset of ∼500,000 TFs, grouped into ∼1,700 orthologous groups (OGs) across ∼3,000 species. The most ancestral prokaryotes encoded multiple TFs. Going by known extant functions of these TFs, they possibly regulated sugar-fermentation metabolism, sensed overall metabolic state and redox, responded to DNA damage or bound metals; many of which are consistent with some reconstructions of ancestral gene pools and physiologies. The number of TFs as well as their superfamily-level diversity, through evolutionary history, matches expectations against genome size derived from extant bacteria, suggesting pre-LUCA diversification of TF sequence families. Emergence of new TFs along the phylogeny largely followed a smooth cumulative distribution curve, suggesting steady innovation, early in prokaryote evolution, in contrast to eukaryotes, in which a majority of TF families emerged in a burst manner at the ancestors of multicellular lineages. Gains of TFs late in prokaryotic evolution predominantly featured recycling of protein families discovered elsewhere in the prokaryotic tree, consistent with the dominance of horizontal gene transfer in these organisms. We speculate on the difference between the evolutionary trajectory of prokaryotic TF repertoire and compare it with the eukaryotic TF repertoire trajectory. This helps us in understanding the manner in which their TF repertoires have evolved in two different super-kingdoms. The difference between the evolutionary dynamics of TF-repertoires might be due to how complexity is envisioned in these two different kingdoms.

## Introduction

Transactions involving nucleic acids are among the most important functions performed by proteins in a cell. These include DNA compaction, modulation of local structural features such as loop formation and DNA unwinding, processes involved in DNA replication, DNA repair, transcription and translation. Among these, regulation of transcription initiation is often achieved by sequence-specific DNA-binding proteins called transcription factors (TFs), which are encoded in large numbers in many cellular genomes(Browning & Busby, 2004). In our definition of specific TFs, we do not include basal TFs such as bacterial sigma factors or archaeal/eukaryotic transcription initiation factors that form part of the initiating RNA polymerase. And we use the term TFs to specifically refer to regulators of transcription initiation and not those that regulate elongation or termination. Notably, the number of TF-encoding genes scales super-linearly with the total number of protein-coding genes in prokaryotes, showing that, in this form of life, genome growth is accompanied by a disproportionately high regulatory load, leading to the suggestion that such an overhead might limit bacterial genome sizes (Ranea et al., 2006; van Nimwegen, 2003). At the other extreme, small bacterial genomes, for example, endosymbionts and parasites, encode for only a few, and sometimes no TFs. No sequence-specific TF is part of any catalogue of minimal genetic requirements of cellular life (Gil et al., 2004). Thus, when and how did TFs evolve and expand?

TFs are characterised by the presence of specific DNA-binding domains, for example, helix-turn-helix or zinc beta ribbon. These domains belong to sequence families that are part of larger protein superfamilies that often include other nucleic acid and protein-binding proteins. Under the assumption that proteins belonging to the same superfamily share a common ancestor, this suggests that TFs and other kinds of nucleic acid-binding proteins might have evolved from each other. It has been suggested that, in some families, sequence-specific DNA-binding TF activity might have evolved from a non-specific DNA-binding and chromosome-shaping protein (Krishnaswamy & Seshasayee, 2025; Visweswariah & Busby, 2015). The evolution of the very catalyst of transcription - the DNA-dependent RNA polymerase (RNAP) - might have proceeded from indiscriminate transcription (called the ’elongation first’ hypothesis) to sequence-specific and highly regulated initiation of transcription in extant lineages (Werner & Grohmann, 2011). Switching specificities from RNA to DNA and vice versa may also occur. For example, several proteins thought to be DNA-binding have also shown RNA-binding properties (Brescia et al., 2004; Macvanin et al., 2012). Even the bacterial RNAP, when bound to a non-cognate metal ion, demonstrates RNA-dependent RNA polymerase activity (Fox et al., 1964; Lazcano et al., 1988).

Once created, TF repertoires in several lineages appear to have expanded primarily by horizontal gene transfer with the possible involvement of gene duplication (Price et al., 2008; Teichmann & Babu, 2004). TFs have been shown to be relatively less conserved when compared to non-TFs, but these trends appear weaker when the conservation of TFs is compared with that of their own regulatory target genes (Ali & Seshasayee, 2020; Madan Babu et al., 2006). The degree of conservation of TFs is correlated with their regulatory scope, or the number of targets they regulate(Ali & Seshasayee, 2020). Many TFs regulate only a small number of genes and are often encoded adjacent to their targets, suggesting that the TF and its target might propagate as a module by horizontal gene transfer; this could also enable a TF to produce low concentrations by coupled transcription and translation to find its targets efficiently (Hershberg & Margalit, 2006; Kolesov et al., 2007). Despite these findings, the evolutionary trajectory of the repertoire of TFs and other nucleic acid-binding proteins across prokaryotes within the framework of a phylogeny remains to be explored. Whereas TFs’ trajectory in eukaryotes has been explored by Dubey et al., 2025. They observe that the TFs’ expansion happened sharply in multi-cellular eukaryotic lineage.

In this study, using a dataset comprising high-quality publicly-available genomes of ∼2,900 bacteria and ∼70 archaea, we present an analysis of the evolutionary histories of TFs. In particular, we ask the following questions: (1) Did the most ancestral prokaryotes code for a transcriptional regulatory network, as TF theoretically seems non-essential? If so, what was the TF repertoire of the most ancestral prokaryotes, and was it different, in terms of numbers and diversity, from extant organisms? (2) How has the TF repertoire changed over the course of prokaryotic evolution? (3) How does the dynamics of TF repertoire evolution in prokaryotes compare with that in eukaryotes?

## Methodology

### Genome and proteome data collections

2,945 complete and species non-redundant prokaryotic genomes were downloaded from NCBI Genbank. To remove redundancy at the species level, the reference or representative species was given priority over the other strains of the same species. In case no representative or reference species was present, a random strain was chosen. If both representative and reference genomes were available for a species, the reference genome was given priority. The final dataset consisted of 2876 bacteria (2374 representative, 92 reference) and 69 archaea (52 representative).

### Identification of the DNA-binding domain

For the identification of the DNA-binding domain, family-level HMM profiles of two major DNA-binding superfamilies in prokaryotes (HTH and ZnBR) were downloaded from the SupFam database (v1.75) (Gough et al., 2001; Wilson et al., 2009). The HTH motif spanned 13 superfamilies (SSF46955, SSF48295, SSF46785, SSF88659, SSF47454, SSF54909, SSF47598, SSF47724, SSF47413, SSF46689, SSF46774, SSF46894, SSF63592) comprising 495 family-level HMM profiles. Additionally, 15 HTH-Pfam profiles, not represented in the SupFam database, were also added to the HTH set (Dubey et al., 2025). The ZnBR superfamily consisted of a single superfamily (SSF57783), with 13 family-level HMM profiles.

Protein sequences from bacterial and archaeal genomes were searched against the HTH and ZnBR profiles database with a standard cutoff of 10^-3^ using hmmscan from the HMMER suite. Since ZnBR has very low sequence conservation, an additional cut-off of bitscore of 9.96 was applied. This resulted in a total of 735,601 HTH and 71,357 ZnBR proteins being identified.

### Identification of transcription factor orthologous groups

The proteins were clustered into orthologous groups (OG). The clustering was based on the eggNOG database (v5) (Huerta-Cepas et al., 2019). This dataset comprises of 219,930 OG profiles. These profiles were searched against our set of DNA-binding proteins with the standard e-value cutoff of 10^-3^. A total of 678,285 HTH proteins and 64,106 proteins of ZnBR were clubbed into 13,702 and 4,324 OGs, respectively.

To identify TFs, a list of 206 Pfam profiles was taken into consideration, from a previous work (Seshasayee & Luscombe, 2011). These Pfams were searched against the 735,601 HTH and 71,357 ZnBR proteins with score thresholds set at the trusted cutoffs (--cut-tc) pre-defined for each PFAM. A total of 492,499 HTH and 2,549 ZnBR TFs were identified. These proteins mapped to 3288 and 75 OGs for HTH and ZnBR, respectively. An OG was termed a TF-OG if more than two-thirds of proteins belonging to that OG could be identified as TFs as defined using PFAM profiles. The final set of TF-OGs, analysed in this study, comprised 1682 HTH TF-OGs and 73 ZnBR TF-OGs.

### Prokaryotic Species Trees

To understand the evolutionary relationships and trajectory of the TFs, multiple prokaryotic species trees were generated or obtained for our set of organisms: These are *16S rRNA-based tree*: The 16S rRNA sequences were taken from the GTDB database (Parks et al., 2022). For the species with multiple 16S-rRNAs, the sequence with zero ’N’ and the longest length was taken as the representative 16S-rRNA. 2645 out of 2945 organisms were selected for the tree, as 300 organisms had identical 16S-rRNA sequences, which could have resulted in multiple polytomies in the tree.

#### Ribosomal protein-based tree

A total of 6 ribosomal proteins were selected from a set of 400 different marker proteins used in the Web of Life pipeline, and 16S-rRNA was added to supplement this set: rplB, rplA, rpsB, rpsC, rplE, rpsG and 16S-rRNA (Zhu et al., 2019). Using similar filtering as above, for removing identical proteins from each set of ribosomal proteins, the total number of organisms was in the range of 2,758 - 2,887.

#### GTDB-generated trees

The Genome Taxonomy DataBase (GTDB) provides separate trees for bacteria and archaea based on 120 and 53 marker genes, respectively. These trees were pruned to our set of organisms and were directly used for further tree-based analyses.

For the 16S rRNA and ribosomal protein trees, sequences were aligned using Muscle v5 with all possible guide tree permutations (Edgar, 2022). The alignment with maximum column confidence was taken for further analysis. The MSAs were trimmed using trimAL (Capella-Gutiérrez et al., 2009) with the gappyout parameter to reduce the phylogenetic noise in the data.

The trimmed alignment was then used to construct the species tree using iqtree v2 (Minh et al., 2020). The 16S rRNA species tree was made using the GTR+F+I+R10 model, which was obtained via model testing(Hoang et al., 2018; Kalyaanamoorthy et al., 2017). For the concatenated tree, the concatenated MSA was used with a partition file specifying the best model for each protein/DNA (Chernomor et al., 2016). This was done to allow for distinct models of evolution for different proteins/DNA.

Three independent trees were generated for both 16S-rRNA and ribosomal proteins. Out of the three, two for each set were taken for further analyses, based on topological similarity to each other. The trees were rooted via two methods: 1) Midpoint root and 2) Minimum ancestral deviation (MAD) (Tria et al., 2017). Both methodologies produced nearly identical trees.

### Ancestral State Reconstruction

The ancestral states of OGs were reconstructed using a two-state ancestral reconstruction for presence/absence. The ace function of the ape package was used for each OG with two underlying models: 1) equal rate (ER), 2) all rates different (ARD). The model with the highest log likelihood was chosen (Paradis et al., 2004). The presence of the OG on a node was defined by having a probability of presence greater than 0.75.

### First emergence of OGs

The tree was divided into separate paths, with each path starting from the root and leading to an extant tip. For each root-to-tip path, the node showing the first presence of an OG was defined as the node of first gain for that OG (‘*path-level discovery node’*). Some OGs were inferred to have been gained multiple times across different nodes in different paths. To account for these, we also defined an absolute first gain by defining the path-level discovery node with the least branch distance from the root as the *progenitor* node.

### Ancestral prokaryotic transcription repertoire

The ancestral prokaryotic transcription repertoire is defined as the transcription factors present in any of the nodes: the last universal common ancestor (LUCA), the last archaeal common ancestor (LACA), and the last bacterial common ancestor (LBCA). Since multiple sets of trees were generated, for each node, the union of the TF-OGs of 16s and ribosomal tree, and its intersection with relevant GTDB trees, was taken as the final set of TF-OG for these LUCA, LACA, and LBCA nodes.

The GTDB has a separate tree for bacteria and archaea. For the LUCA representative node, the union of GTDB-LACA and GTDB-LBCA was taken as a proxy for the GTDB-LUCA.

### Entropy Calculation

TFs belonged to multiple HTH superfamilies, with the majority of the ancestral OGs belonging to the winged HTH. To understand if the superfamily diversity of TFs distribution in any extant genome or internal node, the normalised Shannon entropy was calculated for each node on the tree. The higher entropy would indicate greater superfamily diversity at a node.

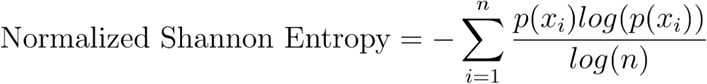

*n, x_i_* is the total number of superfamily and the proportions of TF OGs of *i* family at a node, respectively.

### Simulations and controls

As a control, 100 different simulations, to trace the evolutionary trajectory of the presence/absence of an OG, were run using an in-house R script. The simulation was run with an underlying transition matrix obtained, for each OG, by the ancestral state reconstruction protocol.

A set of random ∼20,000 eggNOGs was also taken into consideration, as proxies for the proteome size.

## Results and Discussion

### Distribution of TFs in prokaryotes

To analyse the evolutionary histories of TFs – defined as non-basal regulators of transcription initiation - in prokaryotes, we assembled a species-level nonredundant dataset of ∼2,900 complete bacterial and ∼70 archaeal genomes. A large majority of DNA-binding proteins in prokaryotes contain either the helix-turn-helix (HTH) or the zinc beta ribbon (ZnBR) motif (Aravind & Koonin, 1999). We searched the genomes in our dataset for proteins containing these motifs. From these sequence searches, we arrived at a dataset of ∼735,000 HTH and ∼70,000 ZnBR proteins. In line with what is known from the literature, HTH is prominent in both bacteria and archaea, whereas ZnBR proteins are found in good numbers primarily in archaea (Aravind & Koonin, 1999). Based on the EggNOG database of orthologs and functional annotations, and a set of DNA-binding domain PFAMs usually found in TFs, we first noted that a relatively small number of ZnBR-containing proteins are TFs, and most such TFs are part of the archaeal basal transcriptional machinery (Supplementary Figure 1a,b). Therefore, we limited our analysis to non-basal or specific TFs containing the HTH motif as the DNA-binding domain.

Among the >700,000 HTH-containing proteins in our dataset, ∼490,000 could be classified as TFs. Based on homology to proteins in the EggNOG database, we grouped these TFs into ∼1,700 orthologous groups (OG). We find that the number of TF OGs is largely proportional to the number of TF proteins per genome, but there is some saturation at higher protein counts, consistent with the idea that multiple members of the same TF sequence family could get clustered together within the same OG (Supplementary Figure 2a). We also took a random sample of ∼20,000 OGs and scanned all the genomes in our dataset for their homologs to arrive at a proxy for the total OG count per genome. In bacteria, which dominate our dataset, the number of TF proteins scales nearly quadratically with total proteome size (exponent = 1.78) (Figure 1a). We see a super-linear relationship between the number of TF OGs and the number of sampled OGs as well (exponent = 1.42) (Figure 1a). Previous studies have suggested that the power law exponent describing the scaling of any subset of proteins with total proteome size might be a reflection of the properties of a family or superfamily as a whole rather than a function within a superfamily (De Lazzari et al., 2017). Our data do not fully support this view for HTH; for, non-TF members of the HTH superfamily in our dataset scale more linearly (exponent = 1.13 at the individual protein count level) with proteome size (exponent = 1.22 at the OG level) (Figure 1b).

**Figure 1:**
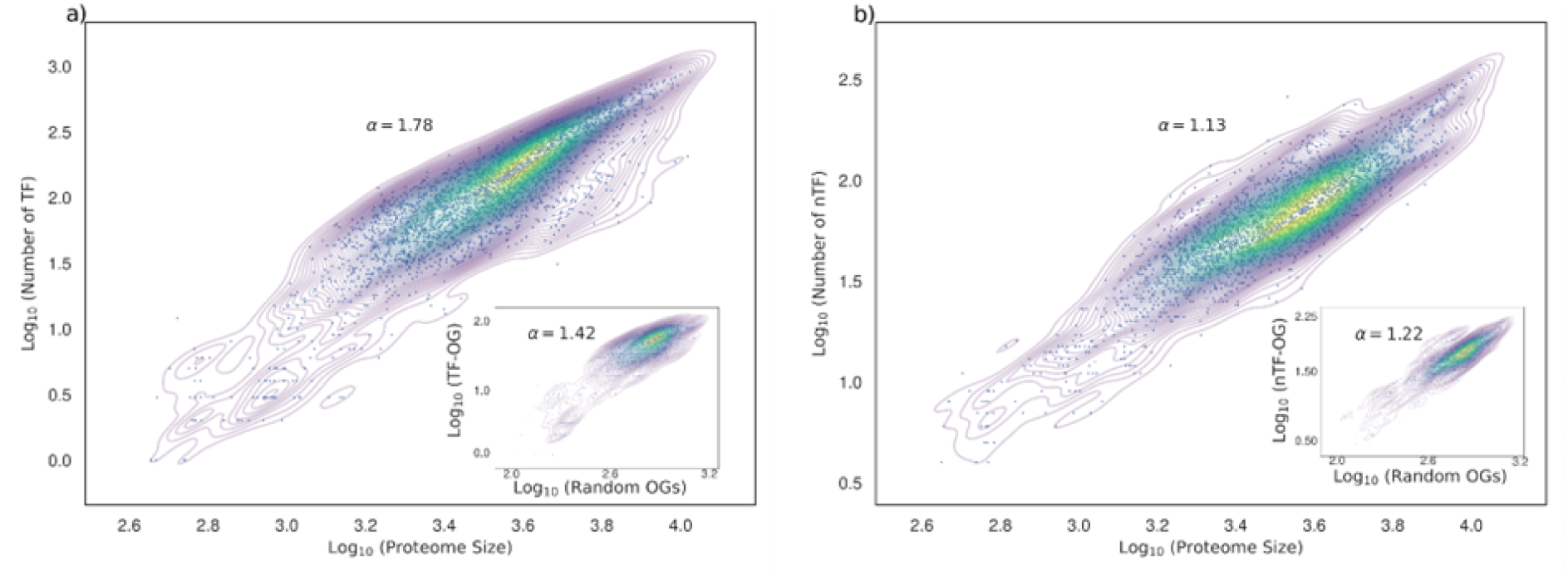
The scaling relation of TF and nTF, versus proteome size within prokaryotes. The scatterplot (log-log) shows TFs (a) scale superlinearly (ɑ=1.78) and nTFs (b) scale linearly (ɑ=1.13) with proteome size across a dataset comprising 2945 prokaryotes. The inset plot depicts the scaling relation at the orthologous group level. The overlaid density plot indicates the density of the data.

### TF repertoire of ancestral prokaryotes

To trace the evolutionary trajectory of the prokaryotic TF repertoire, we assembled a set of phylogenetic trees: two best topologies from (1) 16S rRNA sequence alignments and (2) ribosomal protein alignment concatenation and (3) a pruned tree, one for bacteria and another for archaea, derived from the comprehensive GTDB database, which is based on a set of 120 bacterial and 53 archaeal genes. We rooted the 16S-rRNA and ribosomal protein trees at the midpoint between bacteria and archaea; we also rooted this tree using the minimum ancestor deviation method and found that this produced results nearly identical to mid-point rooting, which we then used for further analysis. We rooted the GTDB-derived tree for the bacterial and archaeal common ancestor as provided by the database.

We used maximum likelihood to trace the presence/absence states of TF OGs across each of the trees. We also did this for all the 20,000 randomly sampled OGs as well as for the non-TF HTH OGs. We first defined the TF repertoire of the most ancient prokaryotes, namely LBCA (Last Bacterial Common Ancestor), LACA (Last Archaeal Common Ancestor) and LUCA (Last Universal Common Ancestor) (Figure 2a). In the absence of eukaryotes in our data, what we call the LUCA might as well be called the Last Prokaryotic Common Ancestor (LPCA); however, in line with the view expressed by Sousa and colleagues that introducing new terms is often unhelpful, we stick to LUCA (Sousa et al., 2016). There is considerable uncertainty in the placement of the LUCA. Therefore, we defined the ancestral repertoire as the set of TFs that show some degree of consensus in their presence across different trees in at least one of the above three ancestral nodes (See Methods for detailed criteria).

**Figure 2:**
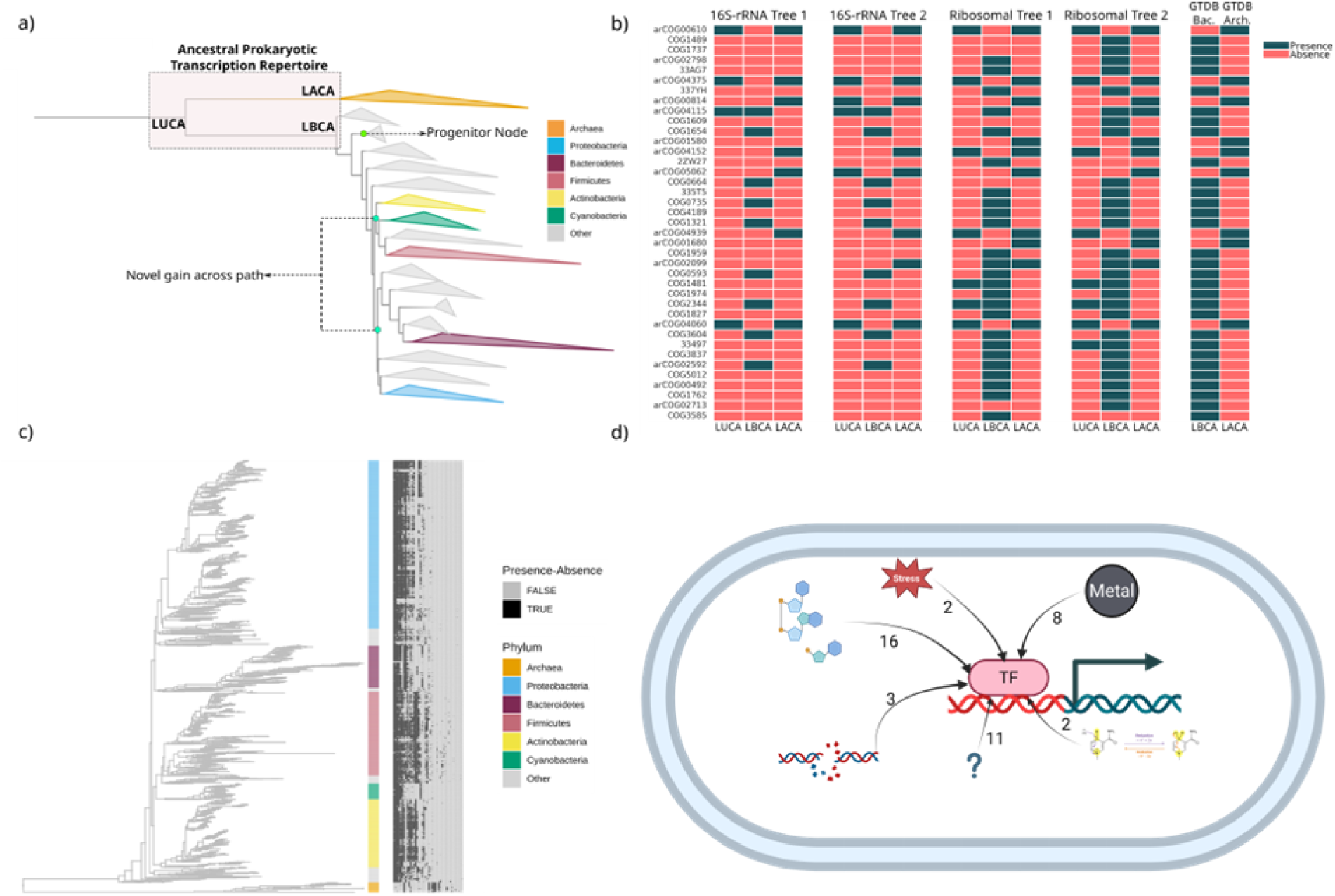
The ancestral prokaryotic transcription repertoire. a) Schematic representation of the ancestral prokaryotic transcription repertoire, depicting the progenitor node and the path-level gain node for an orthologous group (OG). The ancestral prokaryotic transcription repertoire is defined based on the presence of OGs in **LUCA, LACA**, and **LBCA** nodes across multiple phylogenetic trees. (b) Heatmap illustrating the presence or absence of ancestral TF OGs across the three ancestral nodes inferred from different phylogenetic trees. (c) Heatmap illustrates the conservation of ancestral TF OGs across extant species, with phylogenetic relationships overlaid on the 16s species tree. (d) Schematic representation of the ancestral prokaryotic transcription repertoire and its associated functions in accordance with eggnog database

We identified a set of 39 TF OGs in at least one of the three most ancestral nodes (Figure 2b,d). Of these, about 40% (N=17) are conserved in over two-thirds of genomes, and these, owing to the skew in the dataset, are largely bacterial or universal (Figure 2c). Note that six OGs are conserved in at least two-thirds of archaeal genomes, but these are not part of the analysis described below. A highly conserved gene could be called as present in a last common ancestor because it did emerge at and diverge from that node; alternatively, deep or extensive horizontal transfer could also lead to high conservation and therefore trick any ancestral state reconstruction algorithm into inferring its presence in the last common ancestor (Aravind et al., 2005; Weiss et al., 2018). To address this, we visually inspected protein sequence trees for all 17 bacterial ancestral TF OGs and found that only 4 (representing DnaA, LexA, Rex and DtxR) showed obvious separation of protein sequences at the phylum level (Supplementary Figure 3). For LexA, though horizontal transfer between cyanobacteria and alphaproteobacteria has been noted in the literature, the gene tree separates all other major phyla well (Mazón et al., 2004). All these four TF-OGs contain the winged-helix HTH subtype. We were able to align the DNA binding domain of representatives of these four TF-OGs together using structure-guided methods, indicating that they shared a common ancestor (Supplementary Figure 4). The nature of this common ancestor in terms of its DNA binding properties would be interesting to interrogate in future. A majority of the 39 ancient TF-OGs are far less conserved and are often found in the Gram-positive Firmicutes among extant bacteria.

How does the ancestral TF-OG repertoire compare with our understanding of the nature of the physiology of LUCA? There is a great diversity of opinion on the nature of the LUCA. Though there is agreement that this organism was an anaerobe, there are debates on whether it used organic substrates for nutrition (Moody et al., 2024; Weiss et al., 2016). It has also been argued that the LUCA did not even have a DNA genome and instead used RNA (Forterre, 2024). Some authors also suggest that it was a progenote (Di Giulio, 2025). However, consensus analysis of various reconstructions of the genetic arsenal of LUCA suggests that it could have been an organotroph/heterotroph and encoded components of glycolysis/gluconeogenesis (Crapitto et al., 2022; Goldman et al., 2012). This supports our inference that several TFs annotated as regulators of carbohydrate metabolism in extant organisms were found in LUCA/LACA/LBCA. Another analysis of ancestral proteomes has suggested a preponderance of metalloproteins; in line with this, we observe the presence of multiple metal-binding TF-OGs in our set of ancient regulators (Knoll et al., 2016; Zerkle, 2005). In addition to these functions, the TF Rex senses NAD+/NADH ratio through its ancient Rossmann fold, and could have acted as a sensor of overall metabolic or redox status (Aravind et al., 2002; Laurino et al., 2016; Medvedev et al., 2020; Park et al., 2018). DnaA, the initiator of DNA replication in bacteria and an auto-regulator of its own transcription, is an essential protein in many bacteria, with the exception of some cyanobacteria, and is highly conserved (Braun et al., 1985). Finally, the presence of LexA, a regulator of the SOS response in many bacteria as well as of metabolic functions in several lineages, in the LBCA points to the importance of a DNA damage response in this organism (Myka & Marians, 2020; Sánchez-Osuna et al., 2021).

Having established a list of TF-OGs as ancient, we asked if there are any statistical properties that distinguish such an ancient TF repertoire from its extant descendants. For example, did they represent a more limited diversity of sequences than their descendants? In other words, did the sequence or structural diversity of TFs increase with increasing distance from the root of the phylogeny? The 39 ancestral TFs represented 11 out of 13 superfamilies within the HTH motif, the most common being the winged-helix motif, which is in line with observations on extant genomes. We calculated the superfamily-level diversity of TF OGs for each node in the phylogeny using normalised Shannon’s entropy and did not find this to be particularly different for the most ancestral nodes, or for other internal nodes on the phylogenetic tree for that matter, when compared to extant nodes (Figure 3 a,c). Thus, the superfamily diversity of TF DNA binding domains had already been established in the most ancient prokaryotes, indicating that the last common ancestor of the HTH superfamily pre-dated LUCA.

**Figure 3:**
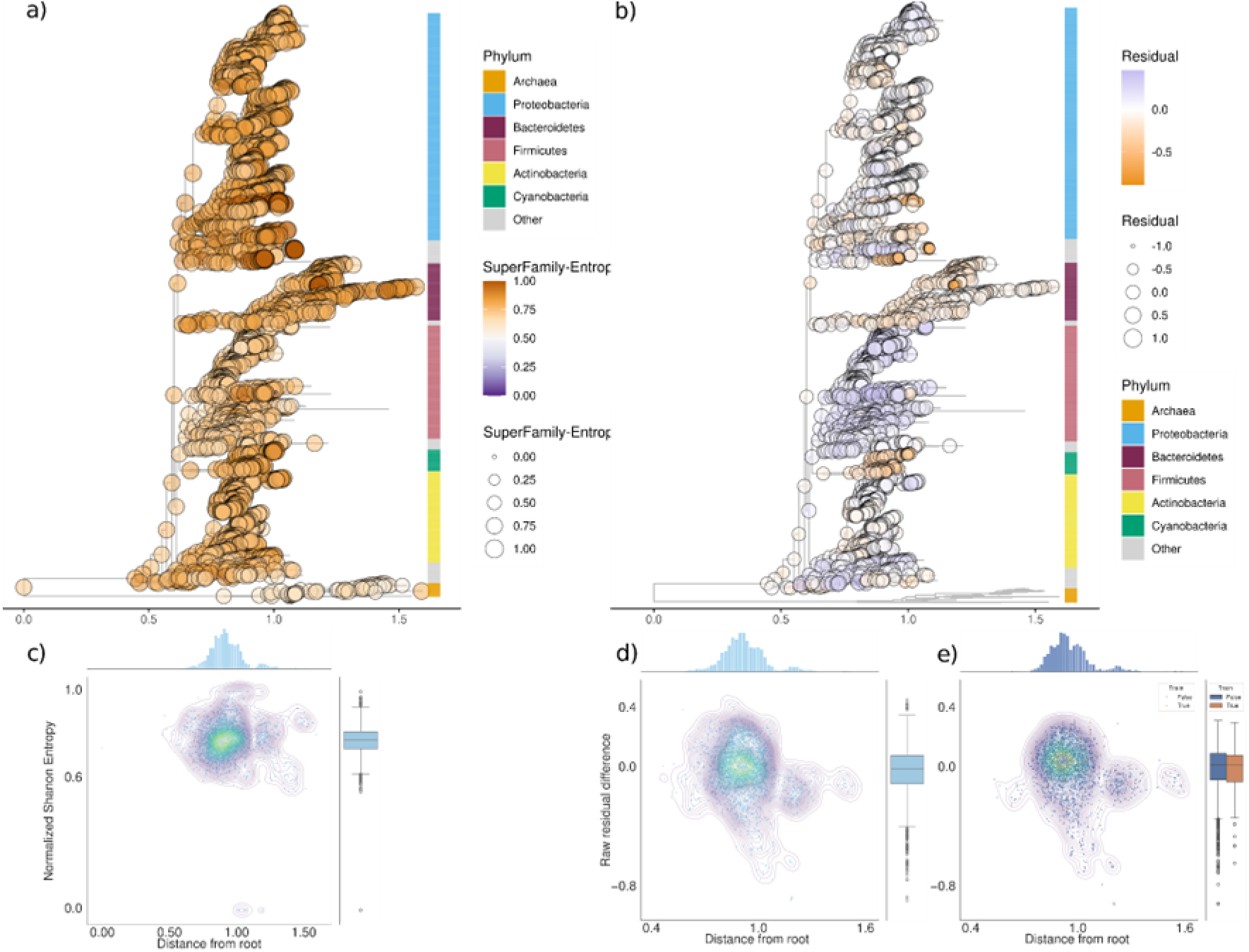
Superfamily entropy and TF-OG superlinear scaling across internal nodes of the prokaryotic species tree. Each node of the 16s species tree represents (a) the normalised Shannon entropy at the transcription factor (TF) superfamily level and (b) the residual difference between the number of TF orthologous groups (TF-OGs) inferred by ancestral state reconstruction and the number predicted by the log-linear regression model based on extant genomes. Scatterplot showing (c) the relationship between normalised entropy and the distance of each node from the root of the species tree and (d,e) the raw residual for internal and extant nodes, based on a super-linear relationship between TF-OGs and proteome size derived from extant genomes, respectively.

We then asked if the number of TF OGs predicted in the most ancestral nodes, as well the other internal nodes, differed significantly from expectations derived from TF-OG vs sampled-OG power law fits for extant genomes. In other words, did the super-linear scaling of TF numbers seen for extant prokaryotic genomes establish early in evolutionary time and remain so across the prokaryotic tree of life? To do this, we used the best-fit power law relationship between TF-OGs and sampled-OGs for a training subset (90%) of extant genomes. From this fit, we calculated residuals for individual genomes for the remaining 10% of extant genomes (test set) and for internal nodes on the phylogenetic tree. We found that the distribution of residuals from this calculation for internal nodes did not differ in any significant manner from that for the test set of extant nodes (Figure 3 b,d,e).

A magisterial review of the structure, function and evolution of the HTH superfamily by Aravind and colleagues - published over 20 years ago - has suggested that certain families within the HTH superfamily are broadly conserved across kingdoms of life and are therefore candidates for presence in the LUCA (Aravind et al., 2005). Modern phylogeny-based ancestral state reconstructions free us from the limitation of using broad conservation as the central measure for inferring the presence of a protein in the most ancestral nodes. Family-level analyses consider large sets of proteins with a diversity of functions as units of evolution. In contrast, the OG-level analysis presented here is aimed towards identifying when particular functions within a broader family might have evolved. This is nevertheless not perfect, and this imperfection is a persistent caveat of studies of this nature. Firstly, boundaries separating OGs within a family can be subjective. Additionally, the nature of sequence comparisons, especially across large evolutionary distances (Price et al., 2007), and the fact that many bacterial TFs belong to a small number of sequence families means that several members of the same family (such as the large LysR family of TFs) and from the same organism could get clustered together within the same OG despite regulating distinct gene functions and presumably possessing different evolutionary histories.

Based on functional importance, it is reasonable to argue that basal TFs, which form a complex with the RNA polymerase and are essential for transcription itself, evolved prior to non-basal TFs of the kind we have studied in this work. Ancestral state reconstructions of the bacterial basal TF sigma70, performed on the trees used in this work, trace it back to the LBCA, whereas those for archaeo-eukaryotic basal TFs date back to the LACA or the LUCA. However, there is nothing to suggest that non-basal TFs emerged later. In fact, the non-basal TF repertoire of the most ancestral prokaryotes was as large and diverse as that expected of any extant genome of similar size. It may well be the case that the diversification of TF sequences initiated before the emergence of these ancient cells. These suggest that basal TFs emerged coeval with- or even later than some with specific TF-OGs. As Aravind et al., 2005 have argued, though it is ‘counter-intuitive’ to propose that basal TFs evolved after specific TFs, it is ‘hardly implausible’ based on conservation-based inferences of the HTH repertoire of ancient cells.

Therefore, the most ancestral prokaryotes probably encoded a transcriptional regulatory network, with TF counts normalised for proteome size and superfamily diversity not particularly different from those in extant genomes.

### Dynamics of TF repertoire across the bacterial tree

Given that the most ancestral prokaryotes and bacteria probably encoded for tens of TFs, what is the evolutionary history, as defined by all root-to-leaf paths along the phylogenetic tree, of all extant TF-OGs? Once discovered, how have TF-OGs been gained or lost over time?

From the ancestral reconstruction of presence/absence states of TF OGs on the phylogenetic tree, we calculated the node at which each TF-OG was first discovered along each root-to-leaf path. Note here that by *discovery* of an OG at a node *N* along a path or lineage, we mean that the OG was gained for the first time at *N* along *that* path; it may or may not have been gained similarly along another path at a point unconnected to the node *N* (Figure 2a). We term such nodes *path-level discovery* nodes. In addition, for TF-OGs discovered along multiple lineages or paths on the phylogenetic tree, we also identified the absolute first point of emergence as that path-level discovery node with the shortest distance from the root; this we call as the ‘*progenitor*’ node for that OG (Figure 2a). We consider both path-level discovery and progenitor nodes as two extreme ends of possibility for the following reason. If OG definitions and phylogenetic state reconstructions were beyond reproach, multiple gains would almost certainly indicate horizontal gene transfers, given the prominence of such events in bacteria. However, given that there are uncertainties in where boundaries between OGs are drawn as well as in tree topologies, there can be a case made that multiple gains of what we define as the same OG might in fact refer to multiple OGs with independent evolutionary histories starting from, say, a common ancestor belonging to the same sequence family of superfamily.

A majority (∼60% in all trees) of all TF-OGs were discovered along more than one path on the tree. We had, in an earlier work, performed a similar calculation for TF DNA-binding domain OGs in eukaryotes, in which about only about a third or less were gained more than once (Dubey et al., 2025). This difference between prokaryotes and eukaryotes may be due to the relative dominance of horizontal gene transfer in the former, which can permit a TF-OG discovered by one lineage to be transferred to another, subject to the caveat mentioned in the previous paragraph.

In eukaryotes, large expansions of TF repertoires occurred at ancestral nodes in multi-cellular lineages in plants and animals. This, when displayed as a cumulative distribution function (CDF) of the proportion of all TF-OGs discovered against the branch length from the root shows an early sharp rise, corresponding to plant lineages, followed by a temporary flattening and then a second sharp increase at animal clades (Figure 4c,d). Non-TFs were gained, on average, earlier than TFs, though there are differences between superfamilies in this property (Dubey et al., 2025). TF-OG gains lag behind species diversification. In contrast, the CDF for bacterial TF-OGs is largely smooth and rises fairly early, with 50% of all TF OGs having been seen on the phylogenetic tree by the time ∼10% of all nodes on the tree evolved (∼35% of all nodes when each path-level discovery was counted as an independent event) (Figure 4a,b,e,f; Supplementary Figure 5-6 for similar results across phylogenies). This implies that the progenitor nodes for over 50% of all prokaryotic TF-OGs were ancient, emerging before and around the branch length range corresponding to diversification at the sub-phylum to family level. A boxplot of the distribution of the number of novel TF-OG gains against nodes binned by branch length from the root shows that the rate of gain of new TF-OGs declined rapidly early in bacterial evolution (Supplementary Figure 7-8; Supplementary Figures 9-12 for similar results for non-TFs and random OGs). Differences between TF-OGs and non-TF or random set of OGs are relatively small and inconsistent between progenitor and path-level discovery analysis, precluding any clear conclusions on differences between TFs and other genes. Taken together, these results suggest that in bacteria, TFs, similar to non-TFs containing the HTH motif as well as a random sampling of OGs, evolved and diversified early, unlike in eukaryotes, where large TF innovations occurred as bursts at specific ancestral nodes in multicellular lineages.

**Figure 4:**
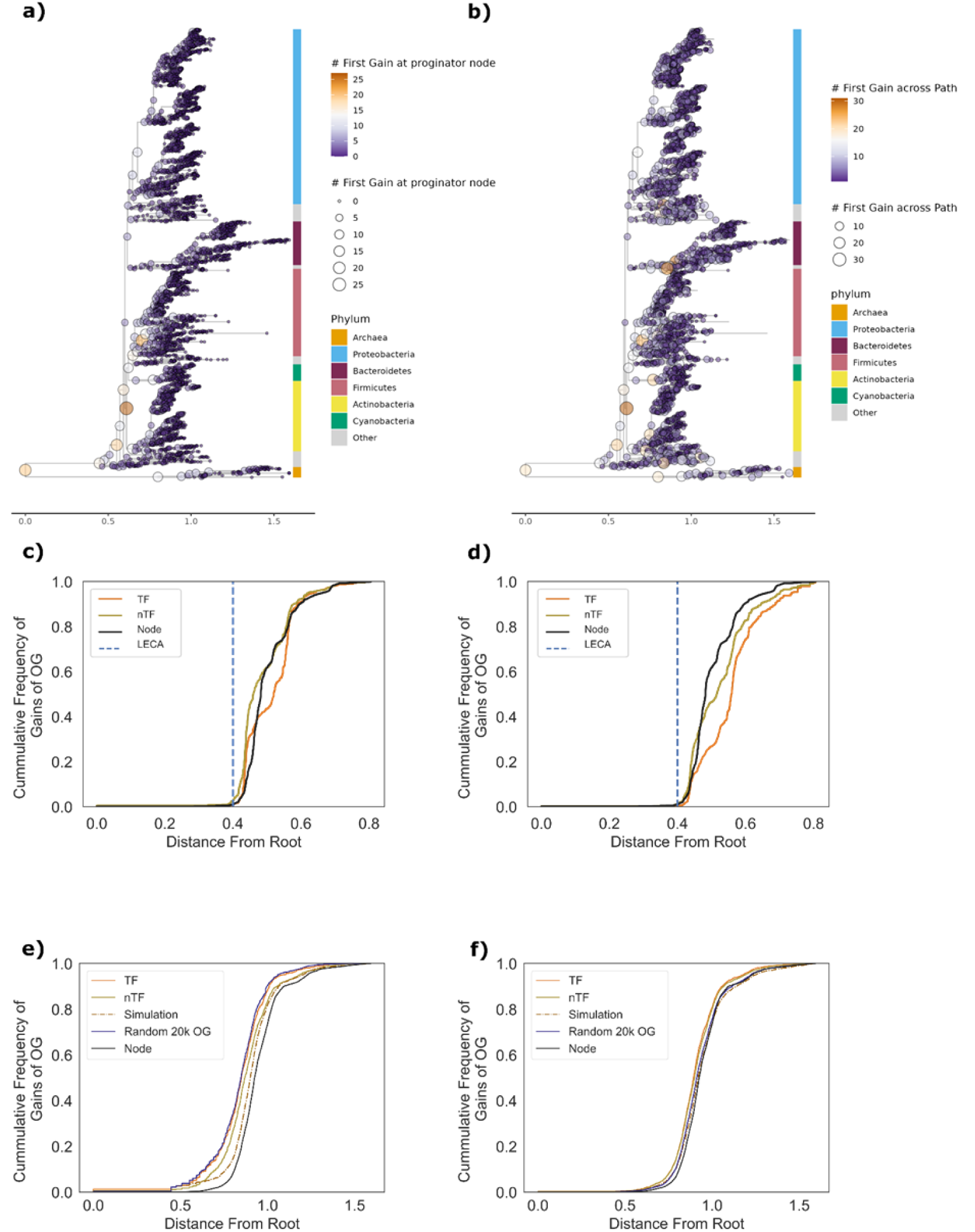
Dynamics of TF-OG repertoire. (a, c, e) TF-OG gains at the progenitor nodes. Panels (a) and (e) show gains within the prokaryotic kingdom, whereas (c) shows gains within the eukaryotic lineage, with a red dot marking the position of the last eukaryotic common ancestor (LECA). (b, d, f) TF-OG gains based on path-level discoveries. Similar to panels (a) and (e), panels (b) and (f) depict gains in prokaryotes, whereas panel (d) shows gains in the eukaryotic lineage, with the dashed blue line indicating the position of LECA. The cumulative distribution functions in c-f show the fraction of TF-OGs (orange) / non-TF OGs (mustard) / random OGs (purple) / simulation TF (dashed brown) and the fraction of nodes in the phylogenetic tree (black) seen at or before the branch length marked in the x-axis.

For each node, we also assembled a list of TF-OGs gained in relation to its immediate known ancestor. This would include not only TF-OGs for which the given node is the progenitor or the path-level discovery node, as well as TF-OGs that were *re*gained following a loss earlier along the path. For any node *N* along the path *P* leading from the root to *N,* we define ‘*innovation ratio*’ as the ratio of the number of TF-OGs gained for the first time at *N* along *P* and that gained at *N* in relation to its immediate ancestor; this value can range between 0 and 100%. In eukaryotes, we had noted that innovation ratio is >80%, i.e., >80% of all gains at a node *N* relative to its immediate ancestor were discoveries along the path. In bacteria, in contrast to eukaryotes, the innovation ratio is around ∼40% across all trees (Supplementary Figure 13). The innovation ratio, when discoveries are defined only by the progenitor node, drops very sharply across the board. Thus, TF-OGs in bacteria are often lost and then *re*gained along the same lineage, in contrast to eukaryotes, where losses along a path are frequently terminal, again consistent with the high probability of horizontal gene transfer events in bacteria.

Therefore, TF-OGs, like many other bacterial genes, including non-TF-OGs sharing the same structural motif as TFs, were discovered early in bacterial evolution and, though lost often, have also been regained and recycled more and more frequently with increasing evolutionary distance from the root, probably by horizontal transfer.

## Conclusions and Perspectives

Transcription is essential for all cellular life. However, its regulation by sequence-specific trans-acting TFs is not. When and how did regulators of transcription evolve? In this study, we have shown that the most ancient prokaryotes probably encoded several TFs whose numbers and diversity were not particularly different from what is seen on average in extant prokaryotes. Most TFs were discovered very early in the phylogeny, starting from the root, but this trend does not uniquely characterise TFs; it applies just as well to non-TFs and to a randomly sampled set of proteins. In contrast, the TF repertoire in eukaryotes has relatively late in evolution, expanding dramatically more than once at particular nodes, especially those within multi-cellular lineages (Figure 5). Bacterial TFs, and other genes, are routinely lost but frequently regained, presumably via horizontal acquisition from other lineages, distant or close; this is again in contrast to eukaryotes, in which TFs once lost along a lineage are rarely regained.

**Figure 5:**
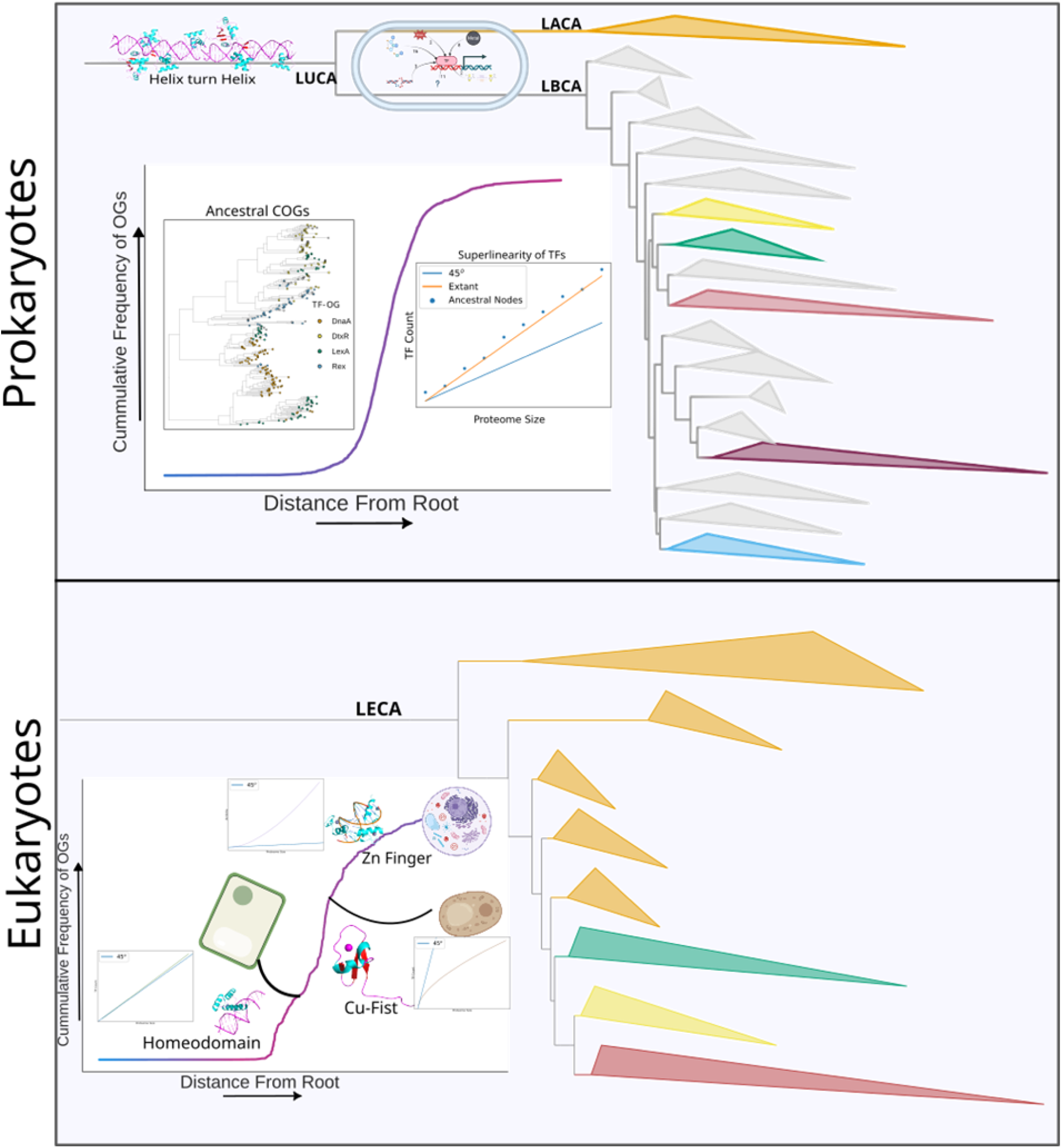
Graphical Summary of the Results. The TF-OGs expanded smoothly over the course of the prokaryotic evolution, whereas within eukaryotes, TF-OGs expanded as burst, which correlated to the expansion of the higher order clades, i.e. Plantae, Fungi, and Animalia. Also, within eukaryotes, we observe bursts correlated to bursts of majorly single superfamilies, in contrast to prokaryotes, where TF-OGs primary helix-turn-helix domain. We also observe that the superlinearity of the TF and proteome has been norm during prokaryotic evolution.

Regulation is something that becomes necessary with increasing complexity, which we define by the need for an organism to partition its genes into partially overlapping sets, each of which is expressed under particular conditions defined in time or space. This need might arise because of cellular economy, i.e., expression of genes not required at a given time-point may be energetically expensive and to such a degree that it impacts fitness; or it could arise because of conflicts between genes by which some genes, when (and where) expressed gratuitously, may antagonise the function of others that are necessary for the organism to fulfil its present demands (Ferenci & Spira, 2007; Lynch & Marinov, 2015; Sorek et al., 2007). Such complexity may increase with an increase in the functional diversity of an organism’s gene repertoire. This might very well be the single most dominant determinant for complexity in bacteria, and more broadly, prokaryotes, across which there is ∼2 orders of magnitude variation in gene content.

In eukaryotes, complexity arises from tissue organisation and multicellularity, and these may find a basis in genome sequence and organisation, but apparently not in proteome size (Hedges et al., 2004). These have evolved in a punctate fashion at particular nodes in the eukaryotic phylogeny. The number of genes in most eukaryotes, across clades, is of the order of 10^4^. However, the number of regulatory macromolecules, TFs in particular, has risen sharply in multi-cellular eukaryotic lineages, in which developmental complexity, as well as the need to manage an excess of the so-called ‘junk’ DNA, has exploded (Dubey et al., 2025; Lynch & Conery, 2003). For example, the ∼3-4 fold difference in the number of protein-coding genes between the yeast *Saccharomyces cerevisiae* and *Homo sapiens* sees as much as a 10-fold increase in the number of TFs (plus a plethora of other regulatory mechanisms), and a massive increase in developmental complexity.

The idea that organism complexity increases monotonically with total gene count across bacteria and prokaryotes, in contrast to the multifactorial definition of eukaryote complexity, is possibly reflected in several aspects of TF evolution in these kingdoms of life. First might be the diversity and distribution of TF types across clades. Eukaryotic TFs belong to several distinct superfamilies, the alpha-helical homeodomains and the beta-stranded Zinc fingers, both of which are abundantly represented among TFs, being two opposite poles. Zinc finger-containing TFs are more common in animalia and the Opisthokonta more generally, but are relatively rare in plants, in which homeodomains are more prevalent. The fact that both homeodomains and zinc fingers can bind DNA and regulate transcription can only be a result of convergent evolution. These suggest independent expansion of particular superfamilies of TFs in different parts of the eukaryotic phylogeny, each of which are very different in terms of their developmental complexity. In contrast, a vast majority of bacterial TFs contain the HTH motif in its DNA-binding domain, independent of phyla, pointing to a single common ancestor possibly preceding the LUCA, which appears to have already coded for a diversity of TFs. Secondly, the number of TFs encoded in a genome as a function of the total proteome size fits a super-linear, nearly-quadratic curve for most bacteria, but varies quite considerably across different clades of eukaryotes, from sub-linear in some fungi through linear to super-linear in animalia. An explanation for the super-linear growth of TFs in bacteria, however, need not invoke any additional notions of complexity beyond the manner in which new genes are added and how new TFs may be assigned for their regulation (Maslov et al., 2009). Finally, the smooth increase in the fraction of TF OGs that had emerged as a function of distance from the root along the bacterial phylogeny, in contrast to a more punctuated pattern observed for TFs in eukaryotes, may also be a function of how complexity might be envisioned in prokaryotes and eukaryotes (Figure 5).

## Data and Code availability

All supplementary datasets are available on Figshare (https://doi.org/10.6084/m9.figshare.31941684), and the code scripts are uploaded to GitHub (https://github.com/IRSINGH27/Early-evolution-of-the-prokaryotic-transcription-factor-repertoire/tree/main).

## Acknowledgments

We thank Inchara Adhikashreni, Ashitha Arun, Tapamoy Bhattacharjee, Suryasarathi D, Sunil Laxman, Sandeep Krishna, Madhumitha Krishnaswamy, Nitish Malhotra, Ganesh Muthu, Mohak Sharda and Pragya Tiwary for discussions on the project and feedback on the manuscript.

