## Supplementary FIgures for "Early evolution of the prokaryotic transcription factor repertoire"

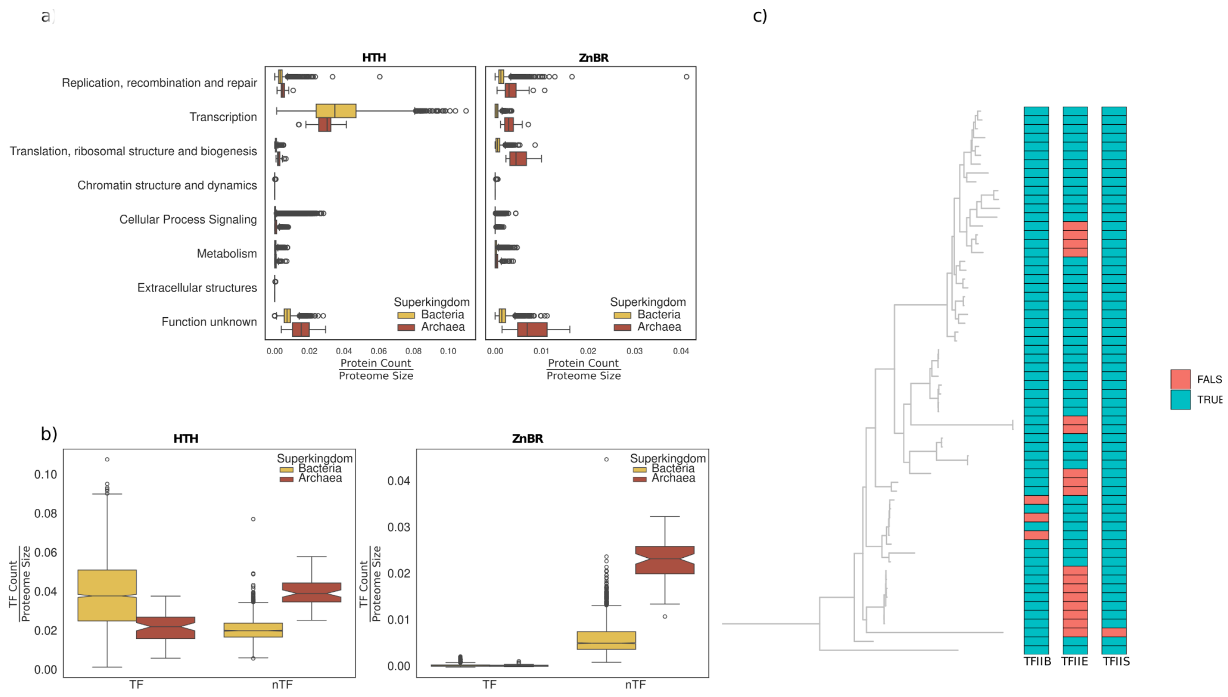


**Supplementary Figure 1. Distribution of HTH and ZnBR proteins.**

(a) Boxplot showing the distribution of proteins across functional categories derived from the COG database. The y-axis represents the COG functional categories, and the x-axis represents the normalised protein count per organism. (b) Similar to (a), but proteins are classified into transcription factors (TFs) and non-transcription factors (nTFs) based on PFAM classification, with the y-axis representing the normalised protein count per organism. (c) Distribution of ZnBR basal transcription factors within the archaeal clade.

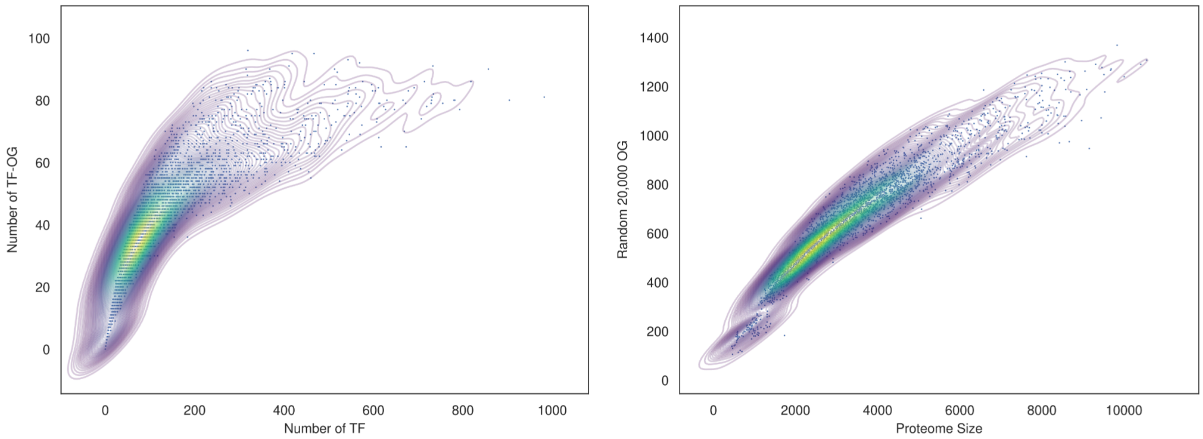


**Supplementary Figure 2. Relationship between proteins and their orthologous groups (OGs).**

(a) Scatterplot showing the relationship between the number of transcription factor orthologous groups (TF-OGs) (y-axis) and the number of transcription factors (TFs) (x-axis). A two-dimensional kernel density estimate is overlaid to illustrate the density distribution and trend of the relationship. (b) Similar to panel (a), but showing the relationship between proteome size and a randomly sampled set of 20,000 OGs, which are used in further analysis as a proxy for proteome-level OG content. Each point in the scatterplot is a single organism.

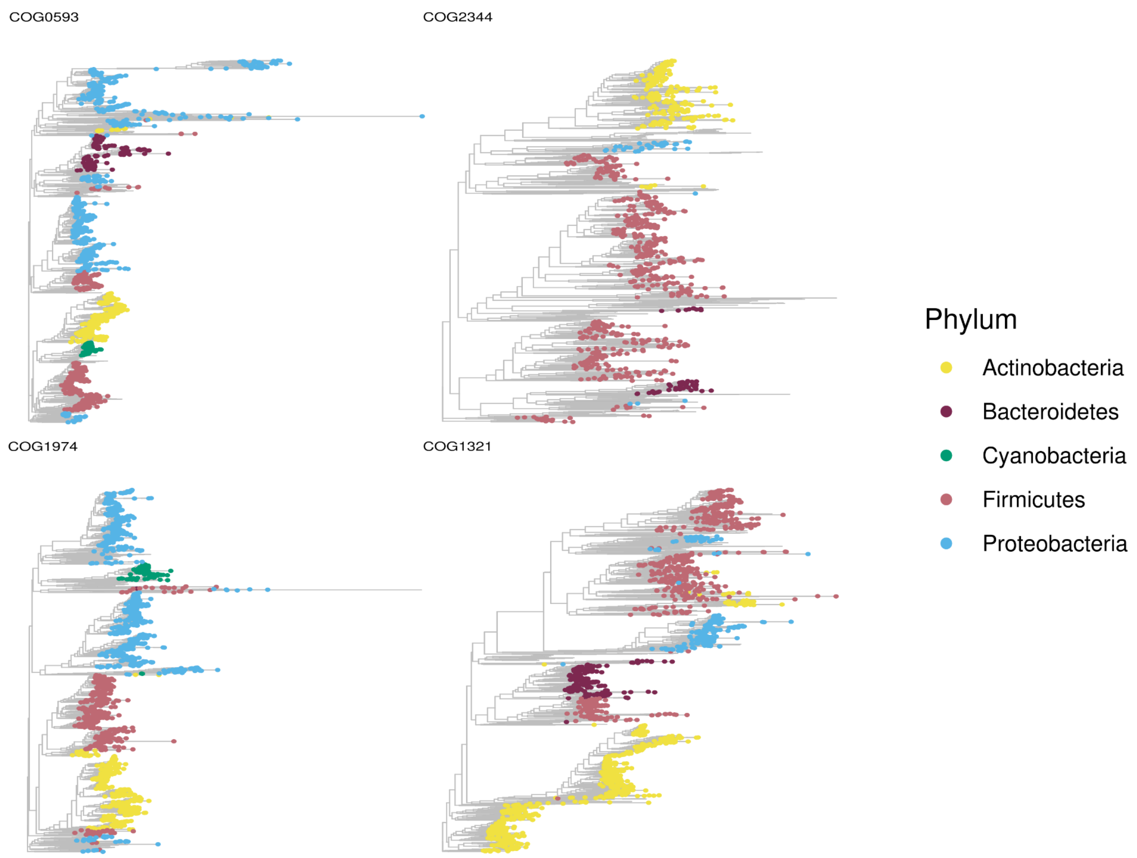


**Supplementary Figure 3. Individual gene trees of 4 ancestral TF-OGs that are conserved in at least ⅔ extant organisms.**

Gene tree of HTH domain with ~ 3000 proteins of dnaA (COG0593), rexR (COG2344), lexA (COG1974), and dtxR (COG1321). The OG, which had more than 3000 proteins were randomly sampled to 3000 proteins. These 4 TF-OGs showed less extent of the HGT at the phylum level, as compared to other ancestral TF-OGs.

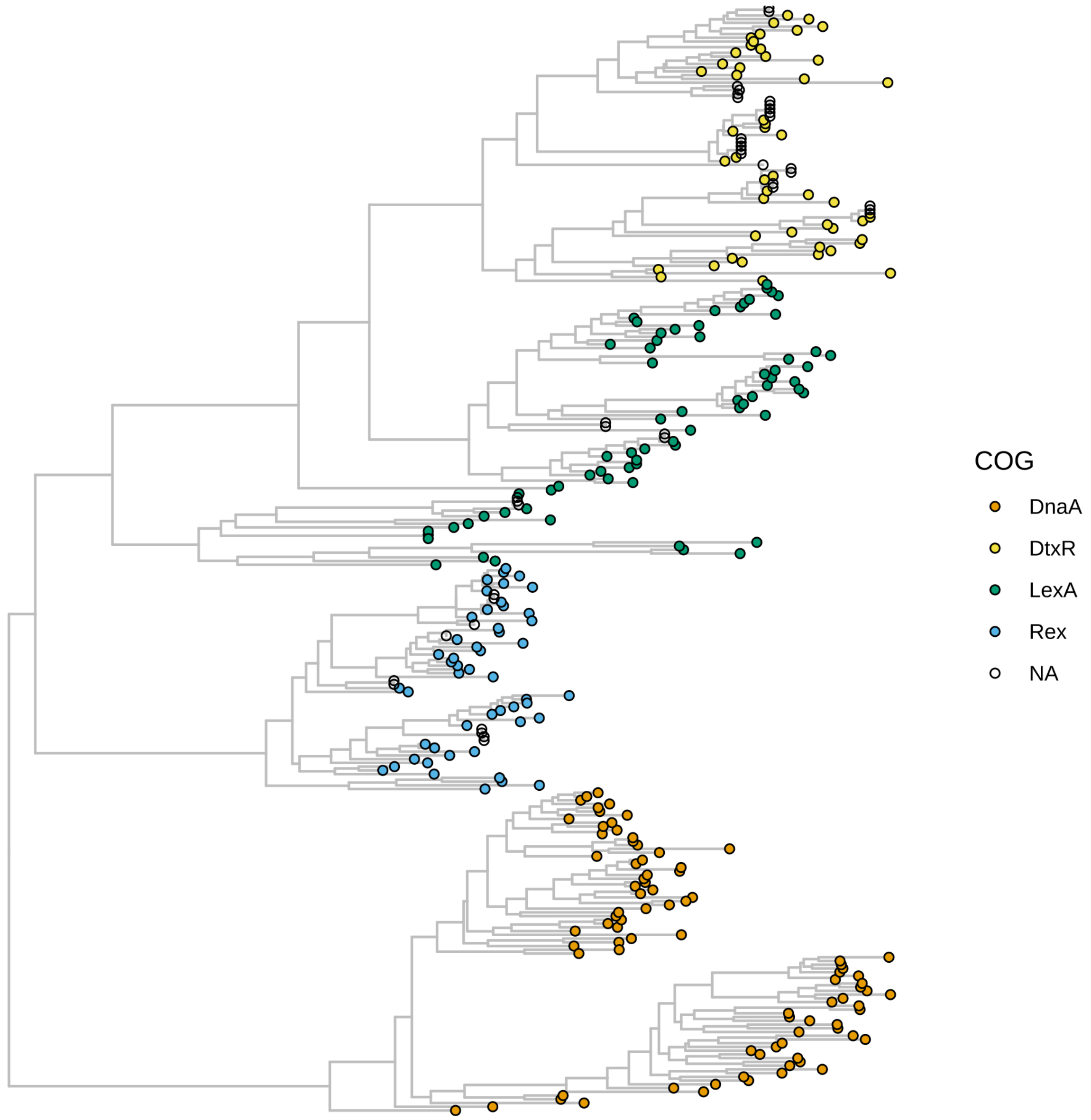


**Supplementary Figure 4. Midpoint rooted gene trees of 4 ancestral TF-OGs.**

The tree consists of 296 sampled down sequences of the HTH domain from 4 ancestral TF-OG sets. The sampling was done via clustering. The midpoint rooting was performed over it, which gave the dnaA clade an outgroup.

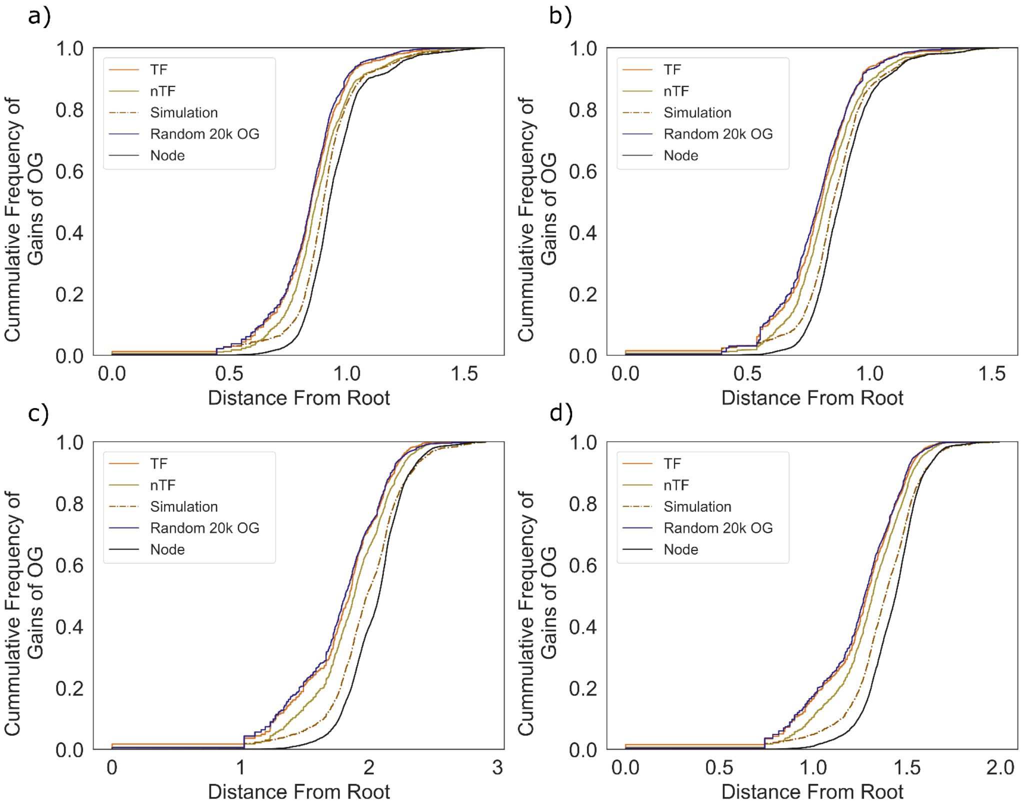


**Supplementary Figure 5. The CDF plot of the TF-OGs over the progenitor node across the tree.**

The x-axis depicts the distance from the root, with the CDF count of the TF-OGs over the progenitor node on the y-axis. Panels a,b,c, and d represent the 16s Tree1, 16s Tree2, Ribosomal Tree1, and Ribosomal Tree3, respectively.

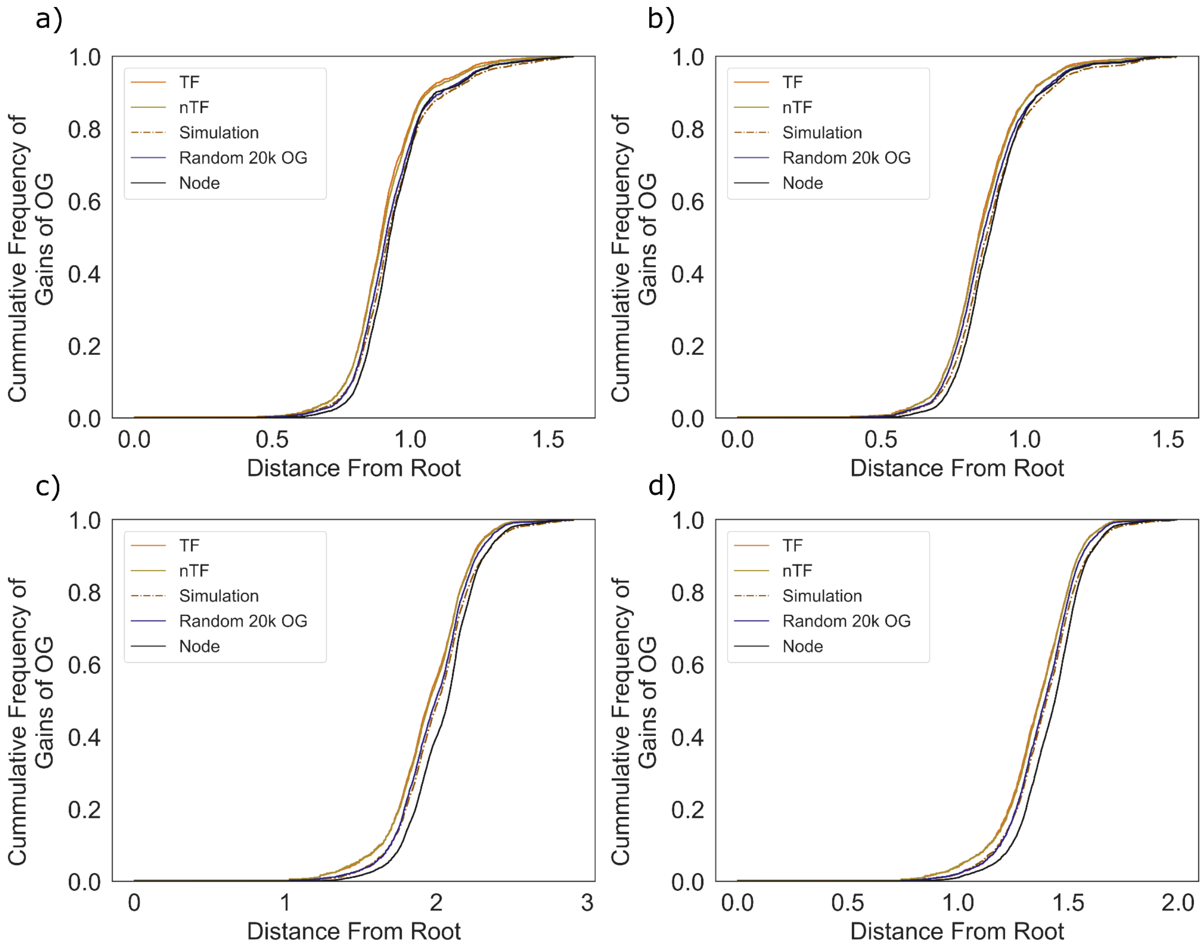


**Supplementary Figure 6. The CDF plot of the TF-OGs novel gains based on path-level discoveries across different trees.**

The x-axis depicts the distance from the root, with the CDF count of the TF-OGs over the first gain node across the evolutionary path on the y-axis. Panels a,b,c, and d represent the 16s Tree1, 16s Tree2, Ribosomal Tree1, and Ribosomal Tree3, respectively.

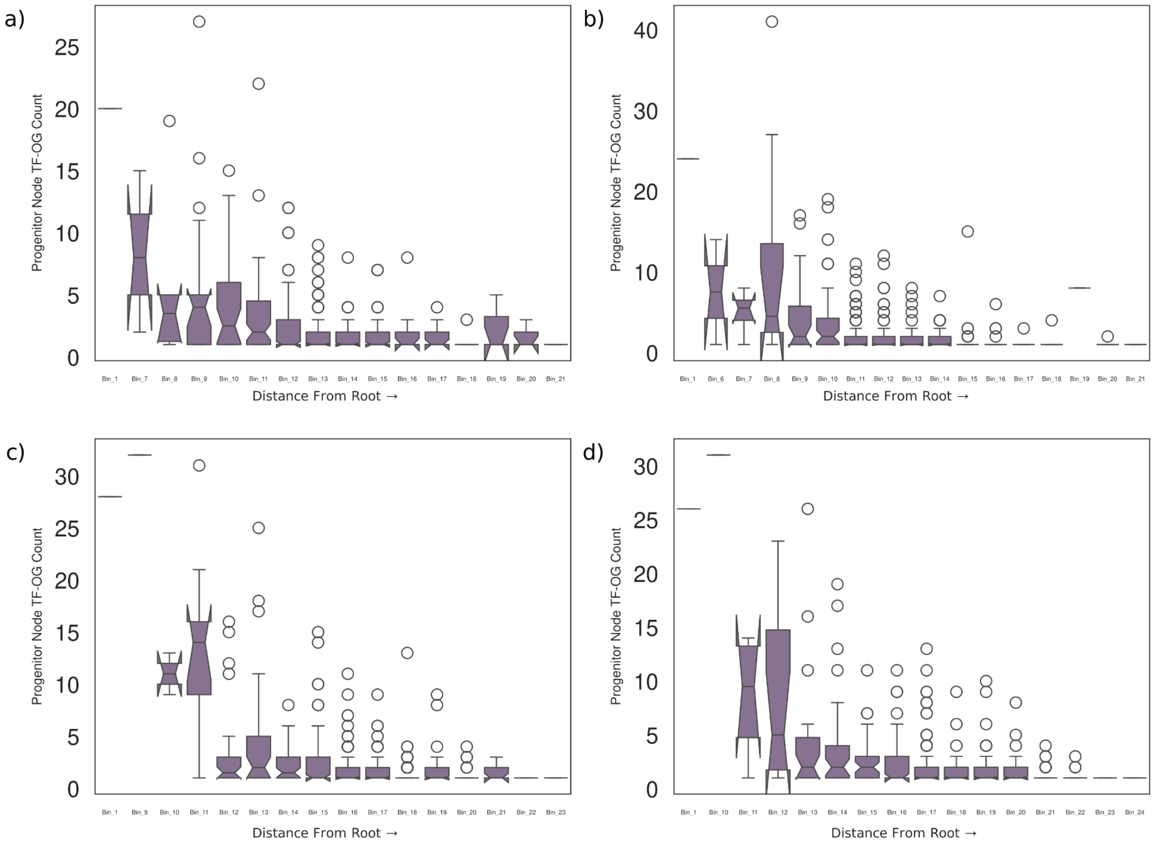


**Supplementary Figure 7. Binned boxplot of the TF-OGs count over the progenitor node.** Each bin is defined by a range of half the standard deviation of the distribution of the distance from the root. The number of nodes in a bin varies as nodes might not be uniformly distributed. (a,b,c,d) represent the trends of the 16s Tree1, 16s Tree2, Ribosomal Tree1, Ribosomal Tree3 respectively.

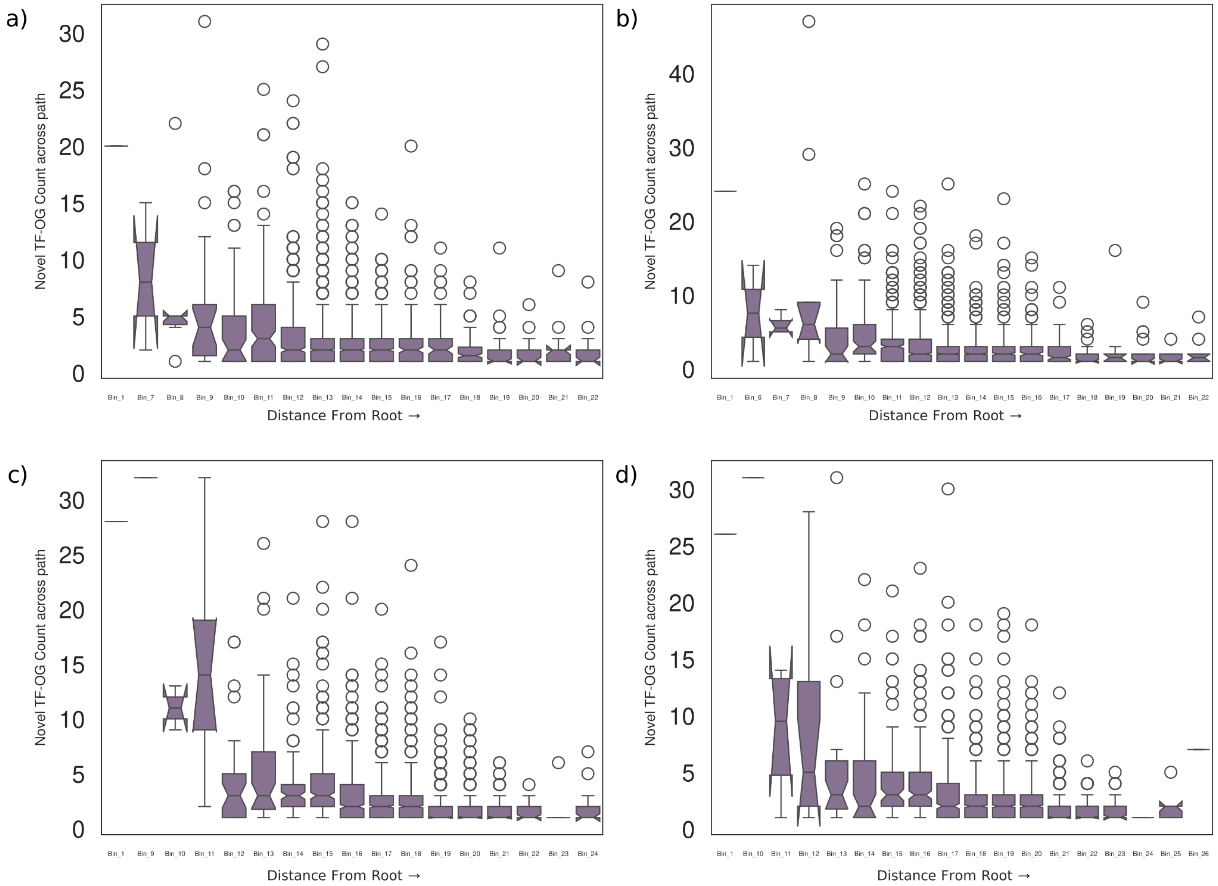


**Supplementary Figure 8. Binned boxplot of the TF-OGs novel gains count over the evolutionary path.**

Each bin is defined by a range of half the standard deviation of the distribution of the distance from the root. The number of nodes in a bin vary as nodes might not be uniformly distributed. (a,b,c,d) represent the trends of the 16s Tree1, 16s Tree2, Ribosomal Tree1, Ribosomal Tree3 respectively.

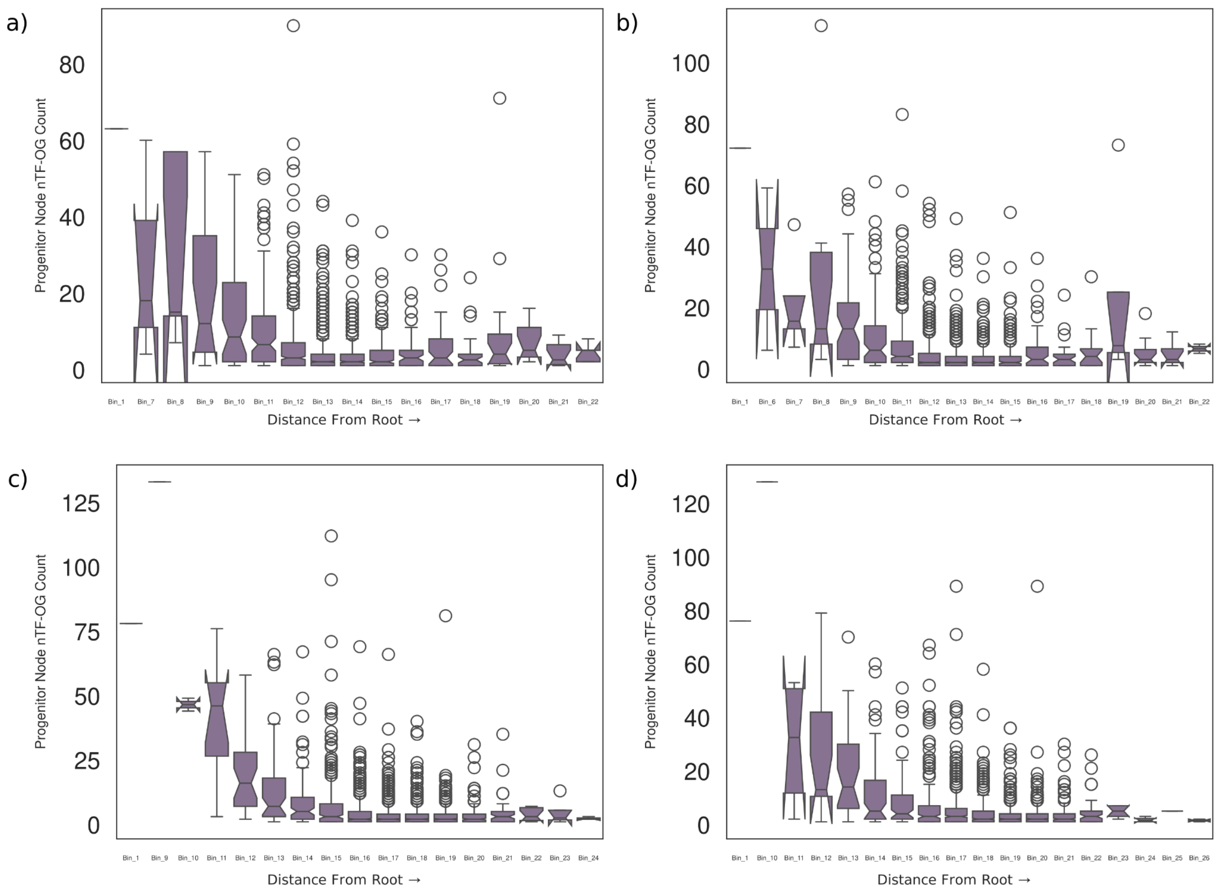


**Supplementary Figure 9. Binned boxplot of the nTF-OGs count over the progenitor node.**

Each bin is defined by a range of half the standard deviation of the distribution of the distance from the root. The number of nodes in a bin varies as nodes might not be uniformly distributed. (a,b,c,d) represent the trends of the 16s Tree1, 16s Tree2, Ribosomal Tree1, Ribosomal Tree3 respectively.

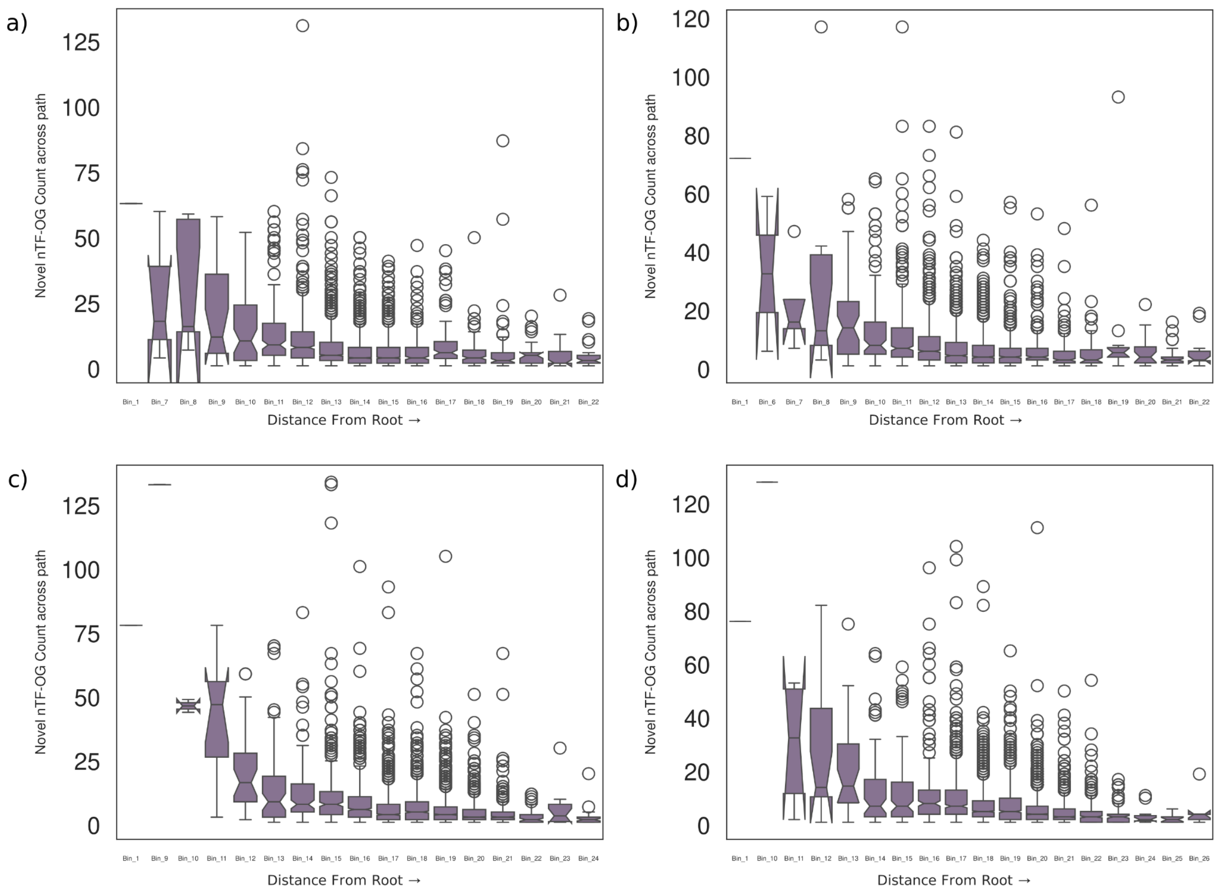


**Supplementary Figure 10. Binned boxplot of the nTF-OGs novel gains count over the evolutionary path.**

Each bin is defined by a range of half the standard deviation of the distribution of the distance from the root. The number of nodes in a bin vary as nodes might not be uniformly distributed. (a,b,c,d) represent the trends of the 16s Tree1, 16s Tree2, Ribosomal Tree1, Ribosomal Tree3 respectively.

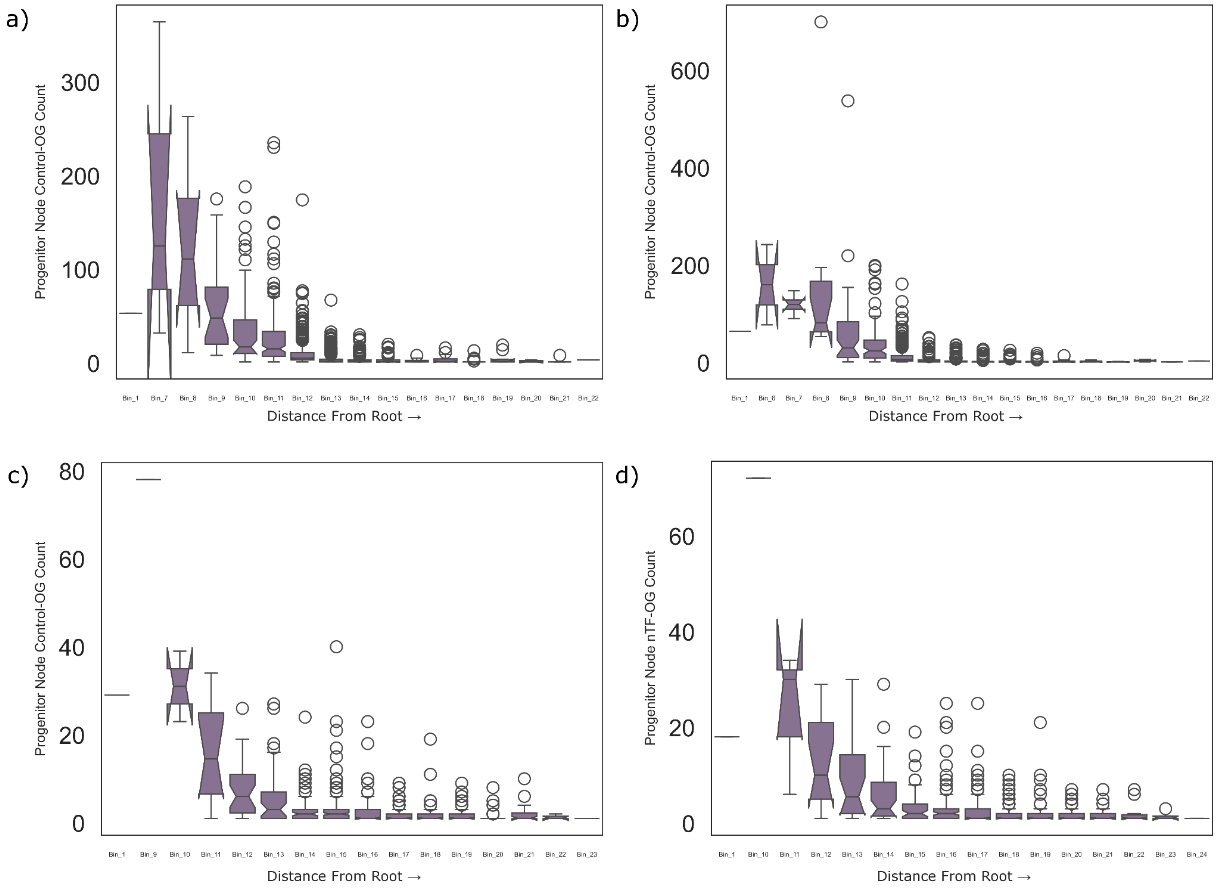


**Supplementary Figure 11. Binned boxplot of the random 20,000 OGs count over the progenitor node.**

Each bin is defined by a range of half the standard deviation of the distribution of the distance from the root. The number of nodes in a bin vary as nodes might not be uniformly distributed. (a,b,c,d) represent the trends of the 16s Tree1, 16s Tree2, Ribosomal Tree1, Ribosomal Tree3 respectively.

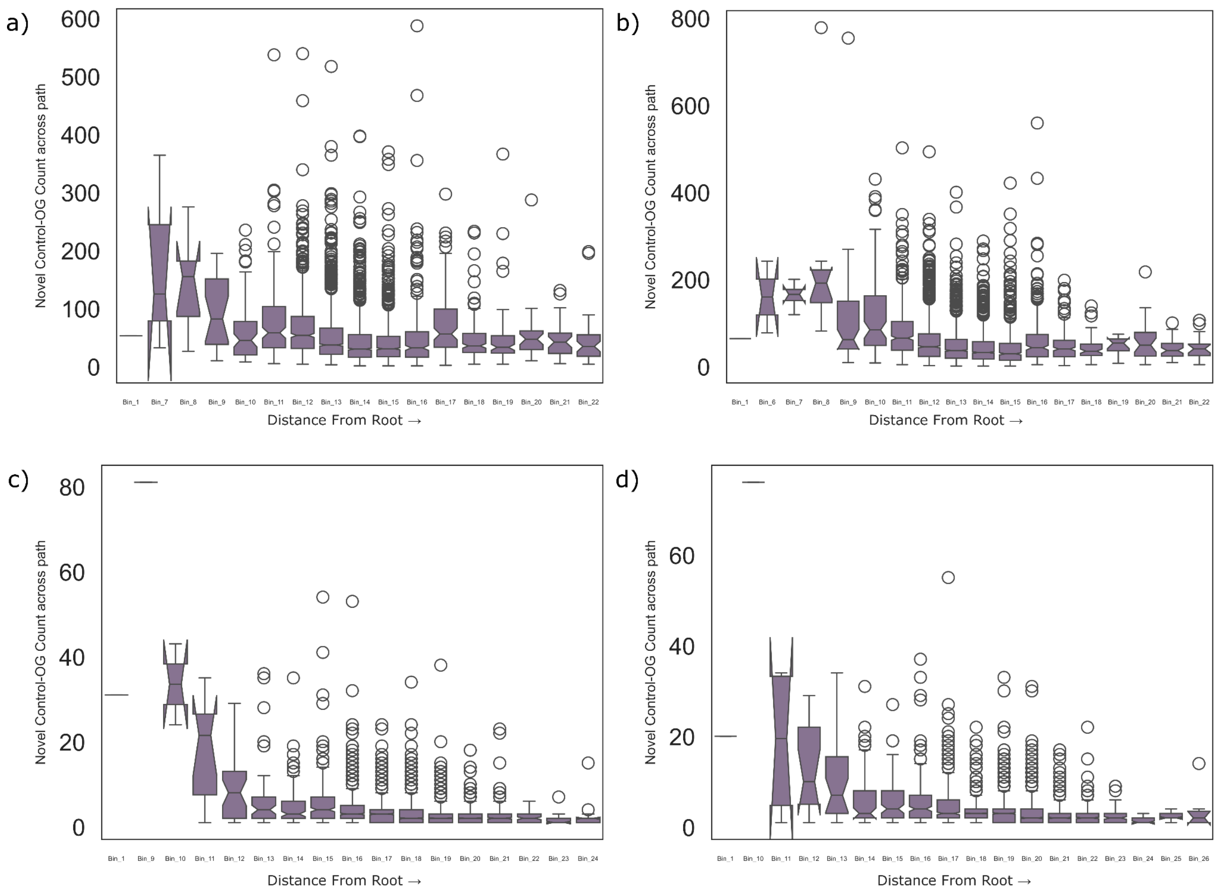


**Supplementary Figure 12. Binned boxplot of the random 20,000 OGs novel gains count over the evolutionary path.**

Each bin is defined by a range of half the standard deviation of the distribution of the distance from the root. The number of nodes in a bin vary as nodes might not be uniformly distributed. (a,b,c,d) represent the trends of the 16s Tree1, 16s Tree2, Ribosomal Tree1, Ribosomal Tree3 respectively.

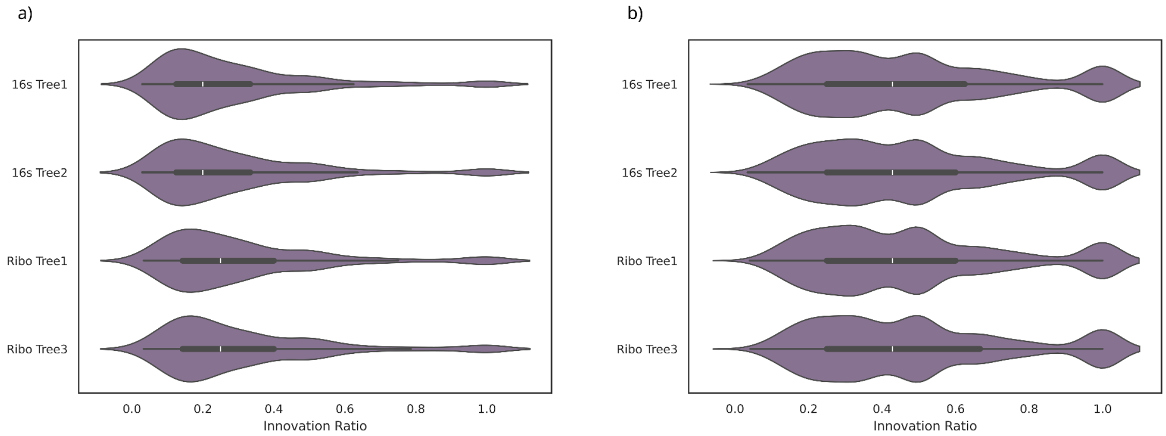


**Supplementary Figure 13. Violin plot of the innovation ratio across the different trees.** Each point in the violin plot depicts the innovation ratio for a node. The x-axis defines the innovation ratio i.e, number of novel gains of the node with respect to the immediate gains from its parent node. The y-axis depicts the different trees. Innovation ratio is calculated based on the progenitor (a) and novel gain across the path (b). Across the tree, we can observe the median innovation ratio to be around 0.2-0.4.
